# DESIGNER DNA HYDROGELS TO STIMULATE 3D CELL INVASION BY ENHANCED RECEPTOR EXPRESSION AND MEMBRANE ENDOCYTOSIS

**DOI:** 10.1101/2021.07.31.454582

**Authors:** Shanka Walia, Vinod Morya, Ankit Gangrade, Supriyo Naskar, Aditya Guduru Teja, Sameer Dalvi, Prabal K Maiti, Chinmay Ghoroi, Dhiraj Bhatia

## Abstract

DNA has emerged as one of the smartest biopolymers to bridge the gap between chemical science and biology to design scaffolds like hydrogels by physical entanglement or chemical bonding with remarkable properties. We present here a completely new application of DNA based hydrogels in terms of their capacity to stimulate membrane endocytosis, leading to enhanced cell spreading and invasion for cells in *ex-vivo* 3D spheroids models. Multiscale simulation studies along with DLS data showed that the hydrogel formation was enhanced at lower temperature and it converts to liquid with increase in temperature. DNA hydrogels induced cell spreading as observed by increase in cellular area by almost two-folds followed by increase in receptor expression, endocytosis and 3D invasion potential of migrating cells. Our first results lay the foundation for upcoming diverse applications of hydrogels to probe and program various cellular and physiological processes that can have lasting applications in stem cells programming and regenerative therapeutics.

## 1. Introduction

Extracellular matrix (ECM) is a 3D natural scaffold produced by different proteins such as collagen, proteoglycans, fibronectin, elastin, and cell-binding glycoproteins^1^. Various components of ECM are connected to form a meshlike intranet between cells, thus providing mechanical strength to the tissues. The ECM is a pool of growth factors and bioactive molecules viz., fibroblast, cytokines. ECM provides adherent substrate/matrix to cells by connecting through cell-binding receptors^2^. ECM controlled various functions of cells, including adhesion, polarity, proliferation, differentiation, migration, gene expression, and apoptosis *in vitro* and *in vivo*^3^. Since ECM is an important component for cell survival, different natural and synthetic matrices mimicking natural ECM structure are considered as potential approaches to probe, program and reprogram cellular behaviours in *ex vivo* and *in vivo* models. Significant research has been done in designing artificial scaffolds that exhibit the properties of ECM such as: enhancing cell growth, mimicking natural ECM, cell maintenance, providing mechanical, biological support; enhance nutritional support and gaseous exchange, metabolic waste excretion^4^. In this context, hydrogels with porous network structure have been studied extensively and employed as an artificial matrix that mimics biological ECM^5^. The unique properties of hydrogels such as, viscoelasticity similar to soft tissues, ability to retain large amount of water, which enable efficient transport of nutrients and gases through the cells^6^. Therefore, *in vitro* cell culturing should be supplied with such an ECM which satisfied both biological as well as chemical requirements of cells. Different polymers used to synthesise hydrogels include polyvinyl alcohol (PVA), polyethylene oxide (PEO), and polyacrylic acid (PAA)^7^. These materials are biocompatible as well as biodegradable, but on hydrolytic biodegradation releases toxic acids that can be harmful for the cells. Also, their hydrophobic nature can lead to nonspecific adsorption of proteins thus resulting in uncontrolled and unwanted cell interactions^7,8^. Thus, to overcome these drawbacks researchers used biological derived materials such as hyaluronic acid, elastin and collagen as cell matrix. However, due to their inherent bioactivity, reduced stiffness, tedious synthesis protocols and variability when synthesised in batches, they were difficult to control and thus can affect immune responses when implanted^9,10^.

Synthetic DNA strands can self-assemble by Watson-Crick base pairing in a mesh-like structure which can swell upon addition of water, called hydrogel^11,12^. DNA hydrogels have showed remarkable properties such as mechanical stability, biocompatibility, hydrophilicity, self-healing ability, precise base pairing leading to uniformity in batches^13,14^. The hydrophilic nature enhances its gelling property on binding with water molecules. Due to these characteristics DNA hydrogels have huge potential in cell biology applications as these can be precisely engineered to form scaffolds that can easily control cell-cell and cell-substrate interactions^3,15^. Such DNA based scaffolds mimicking natural ECM can bridge the gap between 2D *in vitro* cell culture and *in vivo* animal models by reducing the natural complexity. Using the previously established designs of DNA based hydrogels with some modifications, we showed here completely new applications of DNA hydrogels. We have synthesized and characterized DNA hydrogels using single stranded DNA (ssDNA) by self-annealing method **(Scheme 1)**. These hydrogels were used to understand their role in cell attachment, spreading and growth etc. We observe that not only these scaffolds act as cushions for cells attached but also, they stimulate cell spreading, lipid mediated endocytosis ultimately leading to enhanced invasion of cells in 3D matrix. This study will help to improve tissue regeneration applications that are highly dependent on the adhesive property of hydrogels where transplanted cells are mostly anchorage dependent. Most of the hydrogels lack this structurally fine-tuning property. DNA hydrogels with optimized adhesive properties, thus might help in the improved therapeutic response of such transplants.

**Scheme 1:**
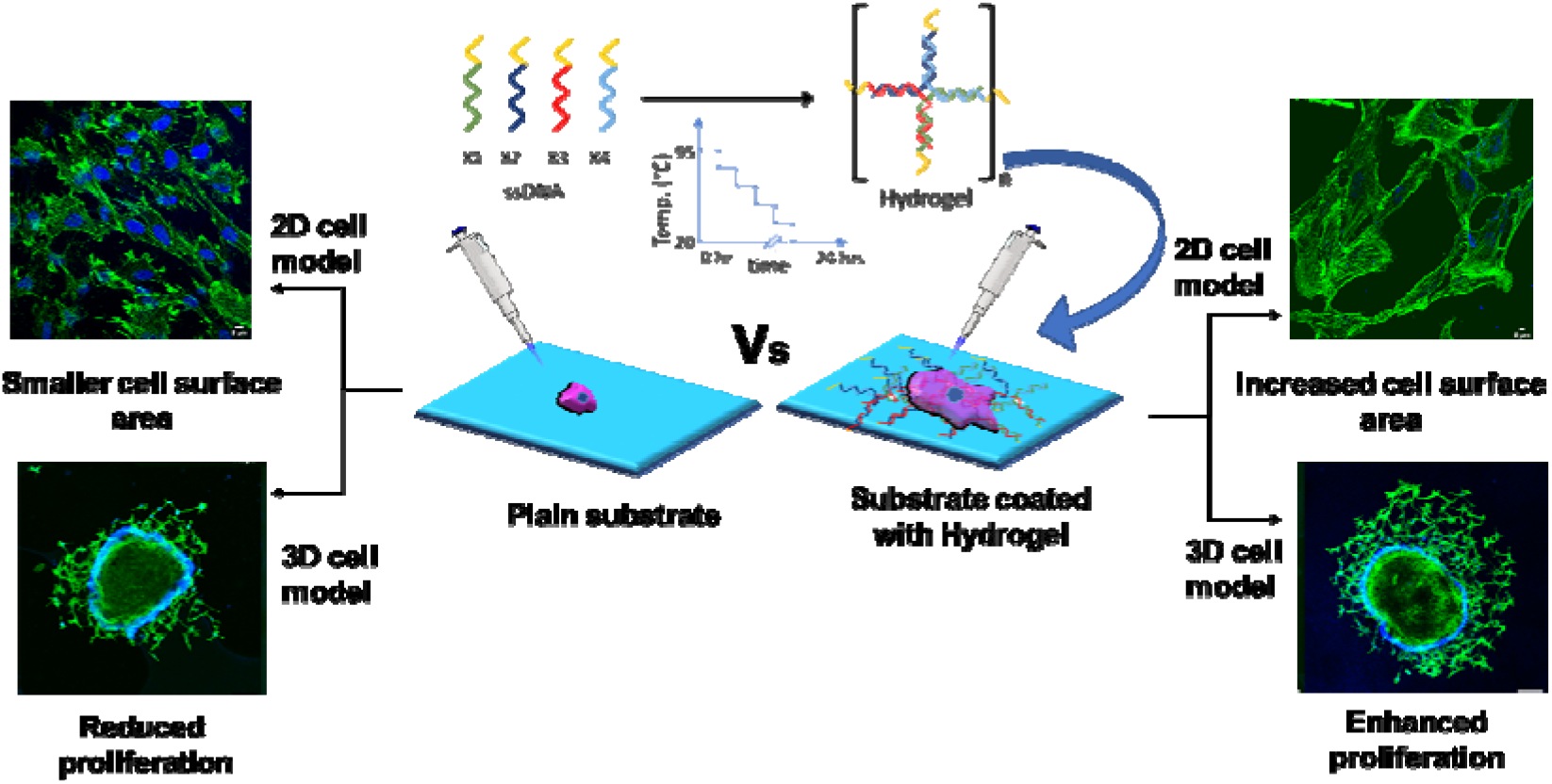
Schematic representation of DNA hydrogels synthesis using ssDNA annealed at 95-20 °C for 8 h and their application in 2D and 3D in vitro cell spreading and growth.

## 2. Results and discussions

### 2.1 Synthesis and characterization of DNA hydrogels

Different types of DNA hydrogels were assembled for motifs modified from previously published results^17^. X-shaped monomer, a four-way junction DNA consisting of X1, X2, X3, X4 oligonucleotide fragments, and resultant hydrogel was named HGX **(Figure 1a and Supplementary information, figure S1)**. Similarly, Y and T-shaped monomers were three-way junction DNA made of Y1, Y2, Y3, T1, T2, T3, and resultant hydrogels named HGY and HGT, respectively **(Figure 1b and 1c)**. We modified the earlier reported^17^ oligonucleotide sequences to ensure higher thermal stability by increasing the G-C content in the sticky ends. Both four-way junction and three-way junction DNA hydrogels were prepared by the self-annealing method **(Figure 1d)**. In order to determine whether we achieve the self-assembled X, Y, T shaped DNA hydrogels, electromobility shift assay (EMSA) was performed to confirm the stepwise assembly of DNA nanostructures **(Figure 1e)**. The mobility of DNA strands decreased as each ssDNA viz., X1, X2, X3, and X4 was consequently added for four-way junction DNA hydrogel, distinct band shift was observed, reflecting the effectiveness of DNA assembly. Similar behaviour was observed for other two types, for three-way junction hydrogels viz. Y1, Y2, and Y3 and T1, T2, and T3. By comparing the mobility of bands in distinct lanes, we concluded that the four-way and three-way junction DNA nanostructures hybridized together to form cross-linked DNA hydrogels, where a band with lower mobility corresponds to a higher molecular weight structure. The temperature dependent formation of higher order structure was also confirmed by DLS where the increase in the size of the structures was observed with corresponding decrease in temperature **(Figure 1f)**.

**Figure 1.**
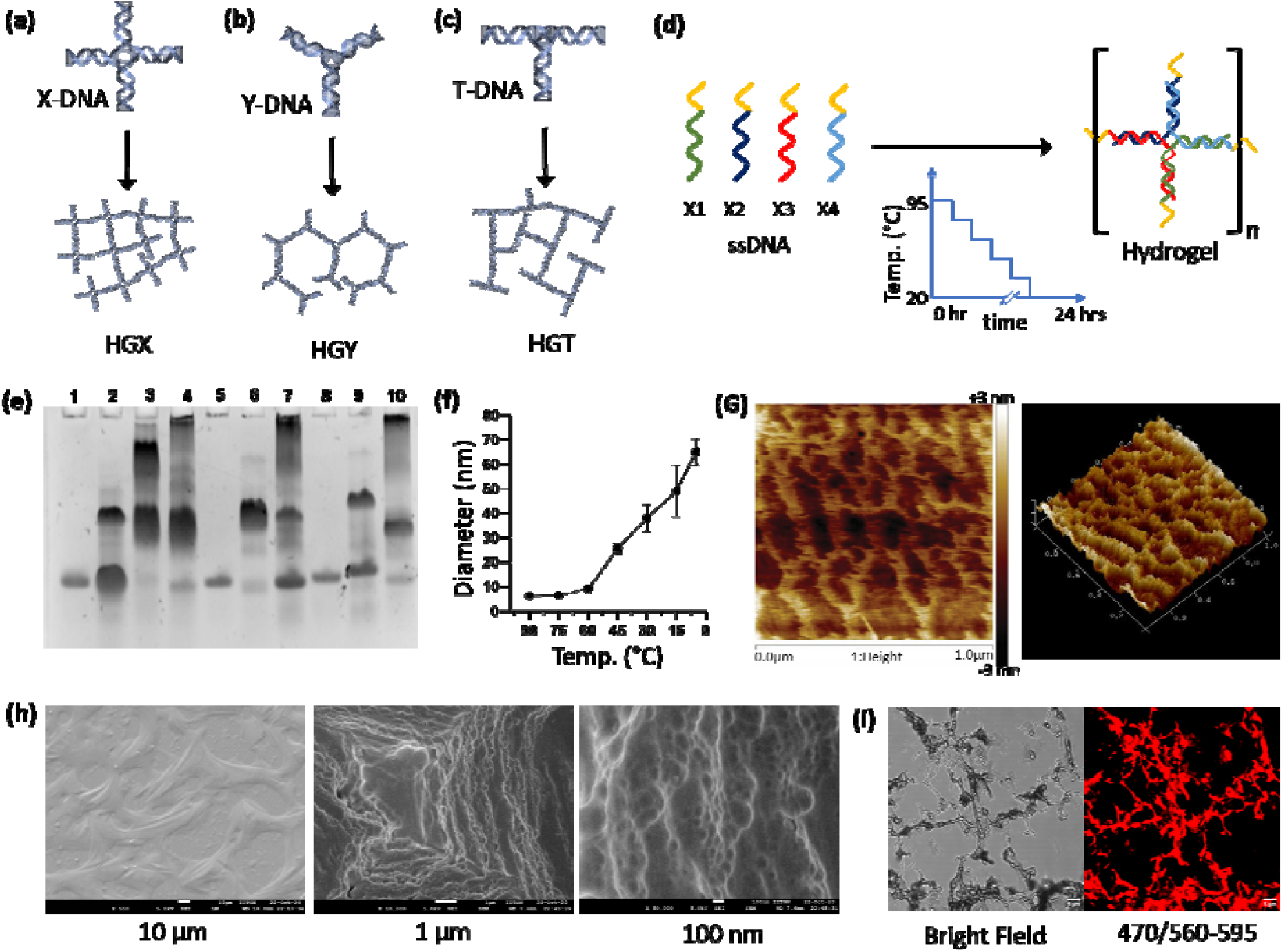
Synthesis and characterization of DNA hydrogels. Schematic representation of (a) X, (b) Y and (c) T-shape monomers and their corresponding hydrogels. (d) Hydrogel formation by self-annealing of ssDNA monomers. (e) Electromobility shift assay to verify self-assembly of the DNA hydrogels: Lane 1: X1; lane 2: X1+X2; lane 3: X1+X2+X3; lane 4: X1+X2+X3+X4 (HGX), lane 5: Y1, lane 6 Y1+Y2; lane 7: Y1+Y2+Y3 (HGY); lane 8: T1, lane 9: T1+T2; and lane 10: T1+T2+T3 (HGT). (f) Temperature-dependent formation of HGX confirming by DLS where increasing diameter showing the building of higher molecular weight structure. (g) AFM images of HGX in 2D and 3D plane (hight gradient bar −3nm to +3nm). (h) FE-SEM micrograph of HGX showing the morphology at 10 µm, 1 µm and 100 nm scale. (i) Confocal images of DOX intercalated in HGX matrix (Scale bar: 5 µm).

The morphological characterization of the superstructures formed was performed using AFM, FE-SEM and confocal microscopy. Atomic force microscopy with tapping mode and force volume measurements were performed on the DNA hydrogels. AFM images also confirmed the porous and network-like structure of hydrogel **(Figure 1g)**. In lyophilized state FE-SEM images showed highly branched and cross-linked structures of DNA hydrogel **(Figure 1h)**. Confocal images obtained in bright-field as well as in visible region showed densely interconnected structures of DNA hydrogel **(Figure 1i)**. These characterization studies clearly demonstrated successful formation of DNA hydrogels by self-assembly process with distinct characteristics of gel like material.

### 2.2. Multiscale Modelling and Molecular Dynamics Simulation

The initial model of all the DNA nanostars (X-shaped, Y-shaped, and T-shaped) was build using Nucleic Acid Builder module of AMBERTOOLs according to the experimental sequence design **(Figure 2a, 2b and 2c**). Model building and simulation details can be found in the materials and methods section. Instantaneous snapshots of each hydrogel system at two extreme temperatures [very high (T* ∼0.775) and very low (T* ∼0.075)] are shown in the inset of **Figure 2d**. From the snapshots, it is evident that at high temperature there are no structural ordering present in the system and the systems behave like unstructured fluid. However, as the temperature is lowered, the nanostars starts to associate with one another through the complementary sticky ends interaction and form a polymeric network. This polymeric network spans through the simulation box, cages the dynamics of the system, and gives its adhesive viscous behaviour. In order to quantify such polymeric network structure, we have calculated the number of connected nanostar pairs. We define that two nanostars to be connected when distance between the sticky ends is <= 0.5 σ_LJ_. We divide the total number of connected bonds with the number of nanostars present in the system to get the fraction of bonds (**Figure 2d**). The fraction of bonds (f) is then fitted with a two-state model as follows: f=1-1/[1-exp{ΔE-TΔS}/T], where T is the temperature, ΔE is the change in enthalpy, and ΔS is the entropy change upon the phase transition. In the two-state model, the system goes from a fully bonded state to a fully unbonded state as a function of temperature. Such a phase transition is mediated by free energy change in the system. When the system forms a polymeric network, it loses a significant amount of entropy which is compensated by the enthalpic gain due attraction between sticky ends. We find that the change in entropic is almost double to that of change in enthalpy as the system undergoes phase transition. The large entropic change originates from the localization of the nanostars in a small confining region when bonded compared to the unbonded state, where they have complete orientation and translation freedom. The contribution of enthalpic change is associated with the interaction of the complementary sticky ends. To understand the microscopic structure of the gel phase, we have computed the radial distribution function (RDF) of the central beads for each nanostar system. The RDF for each system at different temperatures is plotted in (**Figure 2d**). At high temperatures, RDFs do not show any well-defined peak indicating that the nanostar mixture behaves like an unstructured fluid at high T. As the temperature is lowered, well-defined peaks in the RDF emerge, and it becomes more prominent at very low temperatures. These peaks in the RDF imply the growth of the polymeric network at low temperatures. The peak positions depend on the way two nanostars orient and pair with each other. This emergence of the network structures in the gel phase also arrest the overall dynamics of the systems. To probe the dynamics, we have calculated the mean-squared displacement (MSD) of the central bead of each nanostar. The MSD of each system is plotted in **Figure 2f**. At high T, the MSD changes linearly with time, which indicates the dynamics of the structures are diffusive and more like liquid. With the lowering of temperature, the dynamics of each nanostar slowed down due to the formation of the bonds with each other, and MSD vs time deviates from their linear behaviour. For the range of temperatures where MSD show linear behaviour, we have calculated the diffusion constants of nanostars from the slope of the MSD vs t and plotted those as a function of temperature in **Figure S2** of the supplementary information. With decreasing temperature, we find that the diffusion constant of each system is decreasing. When the system forms a percolating network, the dynamics freezes, and the diffusion constant can longer be calculated from the MSD curve. We also find that the diffusion constant of X-shaped nanostar systems is slightly lower compared to Y-shaped and T-shaped nanostar systems. The volume fraction of X-shaped nanostar systems is higher compared to the other two nanostar system, which reduces the available phase space volume to diffuse, and this is reflected in the overall diffusion of the system.

**Figure 2:**
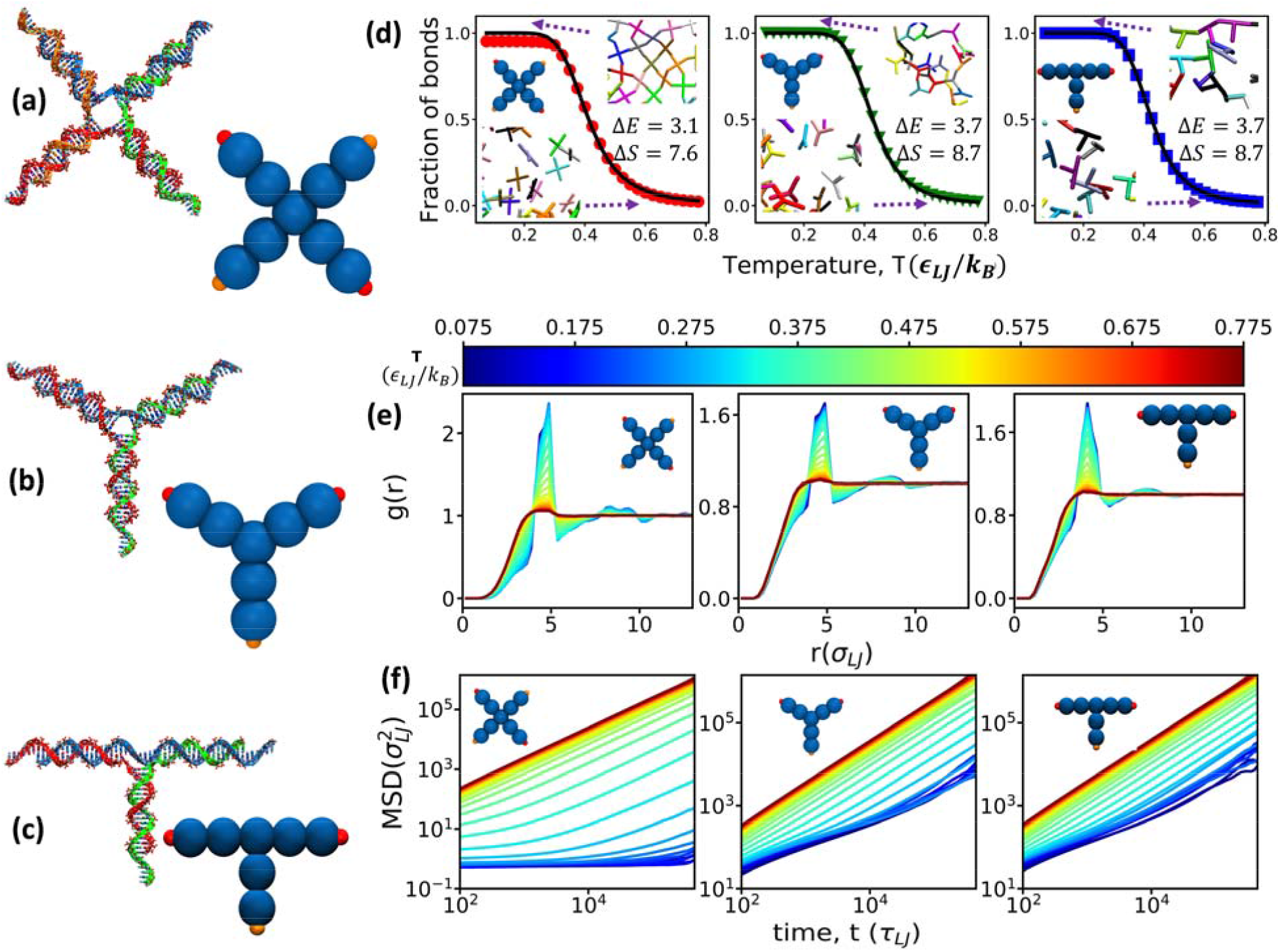
Numerical coarse-grained molecular dynamics simulation study of DNA gelation. (a)-(c) Atomistic and coarse-grained models of each nanostars. (a) X-shaped DNA nanostar. (b) Y-shaped DNA nanostar. (b) T-shaped DNA nanostar. (d) Snapshots each system and fraction of connected nanostar pairs at different temperature. Fraction of bonds Fraction of bonds is determined by calculating the total number of connected nanostar pairs and then divided it by number of nanostars presented in the system. The points represent simulation result, and the straight line is the fitting of two-state model. The fitting parameters are also given in the graph. (e) Radial distribution of function of the central bead of each nanostar system at different temperatures. (f) Mean-squared displacement (MSD) of the central bead of each nanostar system at different temperatures.

### 2.3. DNA hydrogels enhance membrane receptor expression and cell spreading

Adhesion of cells to one another or extracellular matrix (ECM) plays an important role in cell communication and regulation^18–21^. Cell adhesion stimulate signals for cell differentiation, migration and survival of cells. In order to check if DNA hydrogels had any impact on membrane receptors and cellular anchoring via focal adhesions to these scaffolds, we studied the membrane expressions of receptors and cell spread by area calculations. Increase in cell area with and without DNA hydrogels were investigated to check their role in cell spreading. Actin filaments present along the edges of cells resulted in continuous adhesion by distribution and reorganization, thus help in cell spreading^22^.

Actin cytoskeleton when stained and visualize using Phalloidin green, results suggested that on culturing RPE1 cells to these hydrogels viz., HGX, HGY and HGT, cells were nicely adhered and spread on these DNA scaffolds as compared to control sample **(Figure 3 and supplementary information, figure S3)**. From the quantification data it was found that there was significant increase in cell area in presence of DNA hydrogel as compared to control samples **(Figure 3b)**. Also, the actin cytoskeleton was reorganized in cells plated on hydrogels as seen by visible change in their intracellular arrangement (**Figure 3a**). We further noticed that increase in cell area was concentration dependent of DNA hydrogels showing 2 times increase in cell area in case of DNA hydrogel as compared to control sample **(Figure 3 and supplementa information figure S3**). The increase in cell area was comparable to the positive control i.e., poly-L-Lysine. Since all the hydrogels (HGX, HGY and HGT) are showing similar results but the HGX demonstrated best of the spreading results, all the subsequent studies were done with HGX.

**Figure 3:**
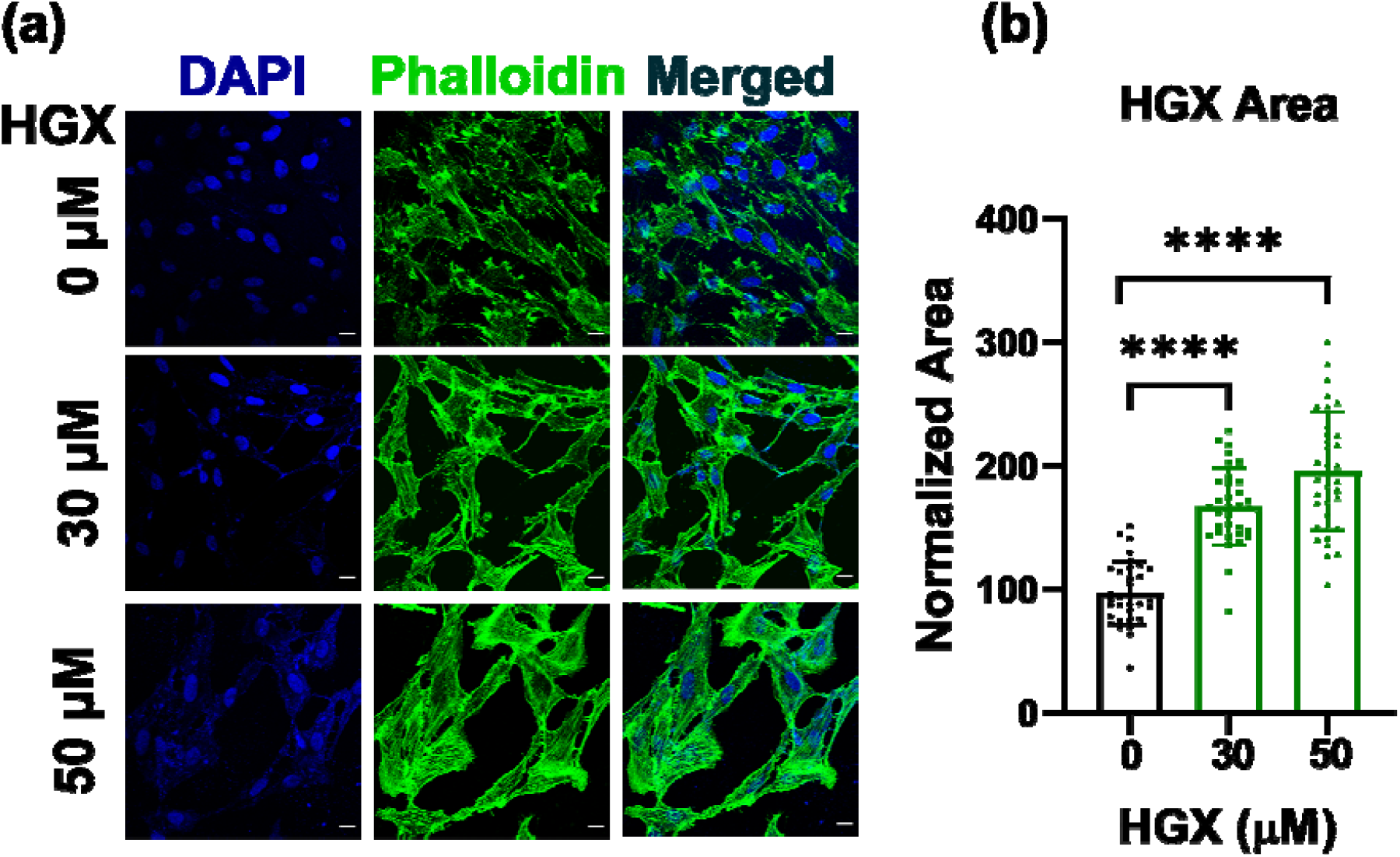
Effect of DNA hydrogels on cellular spreading (a) Confocal microscopic images of RPE1 cells incubated on the carpet of 30 and 50 µM HGX for 24 h, at 37ºC. Scale bar is 10 µm. b) Quantification of normalized cell area w.r.t concentration. **** Indicates statistically significant value of p < 0.0001 (one-way ordinary ANOVA)

Did the increase in cell area on DNA hydrogels affect plasma membrane receptor expression and influence downstream endocytic pathways? To probe these questions, we performed immunofluorescence staining of the plasma membrane fractions of the membrane receptors like CD98, CD147. These are the ectopic receptors present on the plasma membrane and are known to change their expression in response to ECM and disease conditions. We recorded high signal intensity for CD98 and CD147 on HGX coated surface compared to an uncoated surface upon immunofluorescence imaging. To further confirm whether these receptors are overexpressed, or the high fluorescence intensity on HGX coated surface was recorded due to enhanced cell volume, we calculated the flux, the fluorescence intensity per unit area. As shown in **Figures 4c** and **4d**, the flux was also significantly increased, indicating the enhanced expression of CD98 and CD147 on DNA HGX coated surfaces compared to uncoated surfaces.

**Figure 4:**
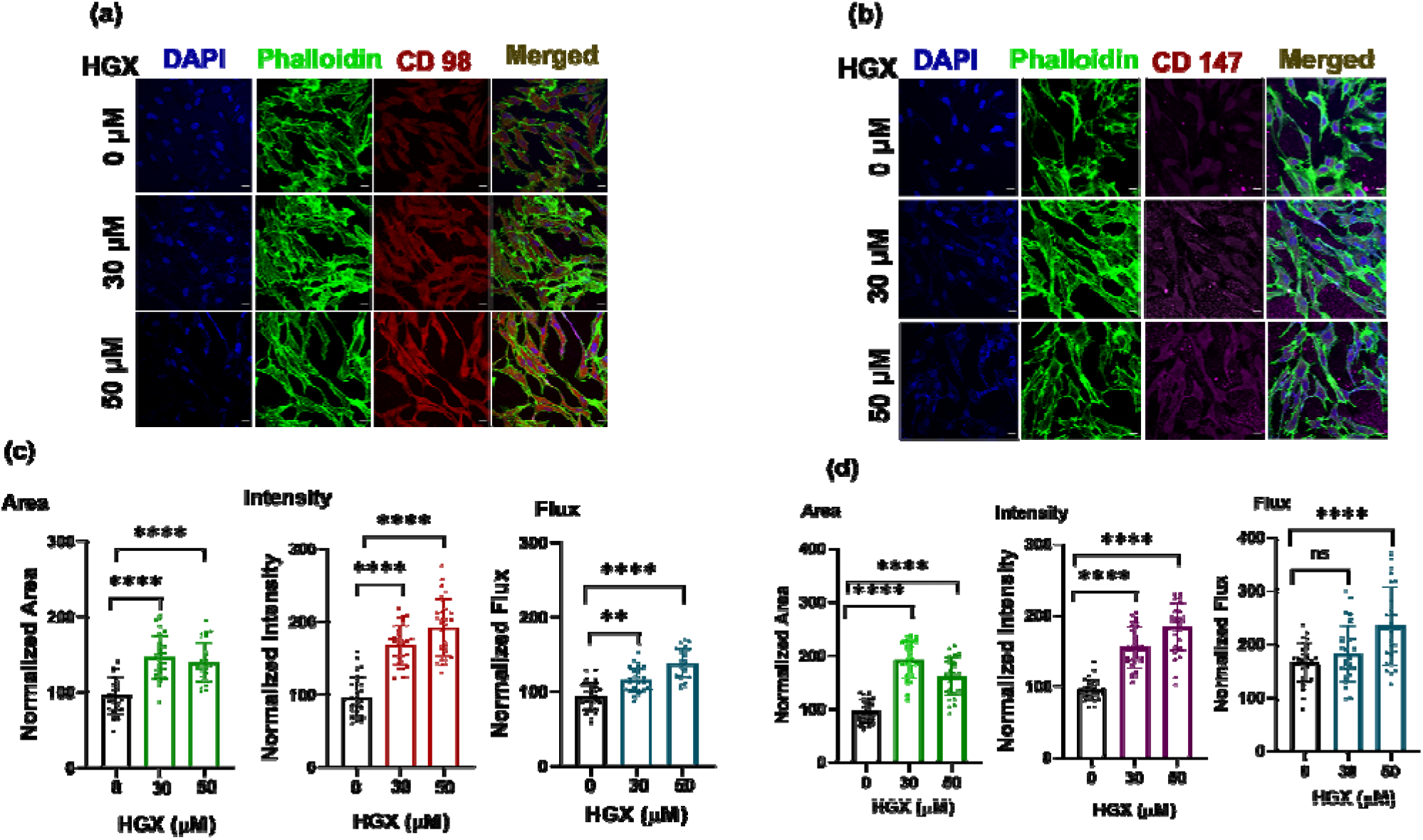
Confocal microscopic images and quantification of Immunofluorescence assay RPE1 cells with and without increasing concentration of HGX incubation (a) with CD98 (red) and (b) with CD147 (red) respectively. In both cases, cellular cytoskeleton was stained with Phalloidin-green (green). (c,d) Cell area, total intensity and the flux (intensity per unit area) for CD98 and CD147 respectively. For quantification, six independent images were captured from each sample, and five cells per image were quantified with the total sum of n=30 using ImageJ software. **** Indicates statistically significant value of p < 0.0001, ** Indicates statistically significant value of p < 0.0073 and and ns indicates no significant difference in the data sets (one-way ordinary ANOVA).

In the case of CD98, cell area was increased up to 1.5 times on treatment with hydrogel, whereas fluorescence intensity and flux were enhanced up to 1.8 and 1.4 times, respectively **(Figure 4a and 4c)**. The change in the area, intensity, and flux in CD147 was ∼1.83, 1.67, and 1.5 times observed in CD147 **(Figure 4b and 4d)**. Increased cell area might have improved the nutritional requirement of cells, causing higher intracellular trafficking of treated proteins. Also, the graph plotted for flux (fluorescence intensity per unit area), showed greater protein levels in the case of HGX as compared to control. However, the change in cellular uptake of endocytic cargos is not depending on the concentration of HGX. It was noticed that there was not much change in cellular uptake of receptor protein w.r.t. to concentration of HGX. Although HGX treatment, resulted in increased surface area of cells. However, increase in concentration of HGX from 30 to 50 µM showed minimal effect on the cell surface area resulting in minimal difference in cellular uptake of treated proteins. This suggested that CD98 and CD147 accumulation was increased in the presence of HGX. Thus, from the study, it was observed that hydrogel increased the cell area, which is related to the enhanced cell migration cell signalling and nutrient uptake.

### 2.4. DNA hydrogels stimulate endocytosis of lipid binding ligands

Endocytosis is a process for internalizing outer material inside the cells and has been implicated in many fundamental cellular functions such as homeostasis, differentiation, signal transmission, cell growth, etc^27^. Endocytosis can take place via multiple routes by either clathrin-mediated or clathrin independents processes. Transferrin (Tf) and galectin-3 (Gal3) are classical ligands used to study the clathrin-mediated and clathrin-independent pathways, respectively.^28^ Cholera toxin B-subunit (CTxB) binds to lipid GM1 on the plasma membrane and builds tubular endocytic tubules which are scissioned by cellular machinery like dynamin or actin or endophilin. These internalized vesicles then mature into early endosomes from where CTxB retrogradely transports to endoplasmic reticulum via plasma membrane and Golgi bodies^29^ We compared the change in cellular internalized fractions of all the above ligands in RPE1 cells seeded over DNA HGX coated and an uncoated glass coverslip. To our observation, the cellular area has significantly increased on the DNA HGX coated surface **(Figure 5B)**. Due to increased cell volume on DNA HGX coated surface, the intracellular uptake of Gal3, Transferrin, and CTxB was also significantly increased, as shown in quantification data **(Figure 5A & 5Ba-c)**. The higher uptake of these ligands is in agreement with increased cell area. The larger cell size may enhance the cell’s nutritional requirement, causing a higher uptake of ligands. In contrast, the fluorescence intensity per unit area, also called flux, was plotted, and surprisingly, we recorded a non-significant difference in the uptake of galectin and transferrin **(Figure 5Bc-d)**. However, a significantly high flux intensity was recorded in CTxB treatment **(Figure 5Be)**. This could be due to lot of protein receptors involved in the endocytosis of Tf and Gal3 while the uptake of CTxB involves mostly lipids on the plasma membrane. In summary, we inferred that the DNA HGX coating enhances the cell growth and area without affecting clathrin and clathrin-independent endocytic pathways but definitely stimulating the uptake of lipid binding cargo like CTxB which drive the uptake of GM1 lipids.

**Figure 5:**
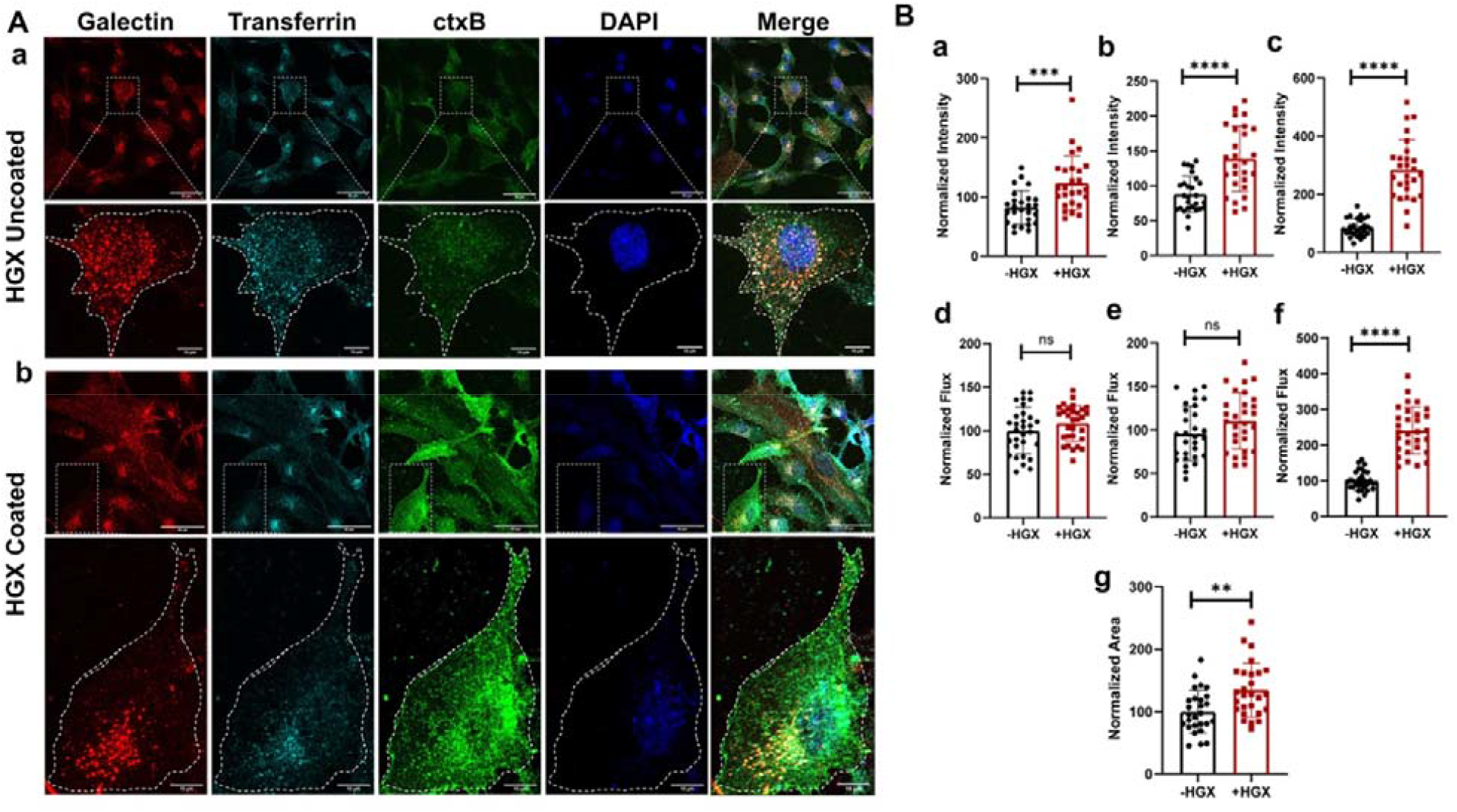
DNA HGX stimulates endocytosis of lipid binding cargos. (A) Confocal images of endocytic probes such as galectin, transferrin, and cholera toxin B (CTxB) internalization by RPE1 cells grown over (a) HGX uncoated and (b) HGX coated glass surface. Scale bar represents 50 µm in zoom out (top row) and 10 µm in zoomed in (bottom row) images of the panel (Aa) and (Ab). (B) Bar diagrams showing fluorescence normalized intensity and normalized flux of galectin (a, d), transferrin (b, e), and CTxB (c, f). (g) Showing representative normalized cell area. For quantification, six independent images were captured from each sample, and five cells per image were quantified with the total sum of n=30 using ImageJ software. The statistic difference was performed using unpaired t-test and indicated by an asterisk (*) in the normalized intensity plot, where **** indicates p < 0.0001; *** indicates p < 0.0003; and ** indicates p = 0.002 and ns indicates no significant difference in the data sets.

### 2.5. DNA hydrogels stimulate 3D cellular invasion

Two-dimensional (2D) cell cultures do not mimic *in vivo* cell growth conditions satisfactorily. 3D spheroid model allows development and interaction of cells with extracellular matrix in three-dimensional environment very much similar to the *in vivo* conditions^30,31^. We thus proceeded to evaluate the 3D cellular invasion behaviour on DNA hydrogels using 3D culture spheroid models. Spherical spheroids made from triple-negative breast cancer cells MDA-MB-231 were transferred to 3D matrix conditions either made of pure collagen, collagen-HGX mix HGX hybrids in different ratios and the spheroids were grown for 24h to study their invasion in 3D. Results suggested that hydrogels (HGX) retained the morphology of cells **(Figure 6)**. Confocal images along with quantification data showed not only successful but enhanced cell migration and adherence in 3D spheroids in presence of hydrogels **(Figure 6a and c)**. It was observed that on incubation with three different concentrations, viz., 25, 50 and 100 µM of HGX, there was differential effect in terms of increased cell invasion at 25 µM of HGX, whereas cell proliferation was enhanced at 50 µM of HGX showing 2 times increase in cell invasion **(Figure 6b)** and an inhibitory effect was observed at 100 µM of HGX.

**Figure 6:**
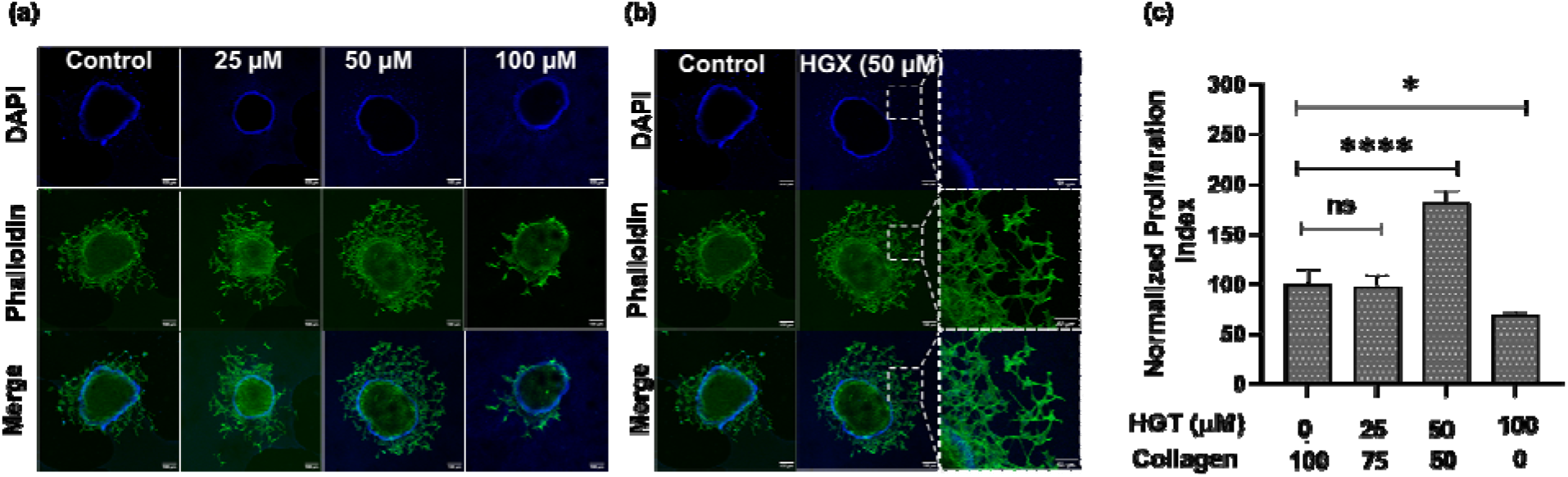
DNA hydrogels stimulate cellular invasion in 3D spheroid model. (a) 3D spheroids grown on different concentrations of HGX for 24 h showed different invasion of cells in 3D matrix as a function of concentration of HGX. Scale bar is 100 µm for all the images. (c) Zoom image of 3D spheroid treated with 50 µM of HGX with 50% collagen showing enhanced cellular invasion from the surface of spheroid in 3D. Scale bar is 50 µm. (c) Quantification data of cell invasion index show higher invasion index for DNA hydrogels upto the concentration of 50 µM. For quantification, three independent images were captured from each sample, and quantified with the total sum of n=3 using ImageJ software. **** Indicates statistically significant value of p < 0.0001, * Indicates statistically significant value of p=0.02 and ns indicates no significant difference in the data sets (one-way ordinary ANOVA).

## 3. Conclusions

We synthesized DNA hydrogels using simple, pre-established protocols using three and four-way junctions via self-assembly method. These DNA hydrogels when characterized showed hierarchical structural properties and striking internal morphology confirming our DNA hydrogels predicted designs. Simulation studies along with DLS data showed the formation of polymeric network structure of hydrogel due to the association of bonds with one another through the sticky ends whereas, these act as unstructured fluid when the temperatures are high enough. This study showed that these hydrogels are thermo responsive hydrogels and can be used as cushions for cellular studies. Cellular studies showed that DNA hydrogels resulted only in enhanced cells adhering to these scaffolds but this further leads to increase in cell area as well as enhanced uptake of endocytic membrane cargos. Cellular area has significantly increased on the DNA HGX coated surface, resulting in increased cell volume, and the increased expression of membrane receptors like CD98, CD147. The larger cell size may enhance the cell’s nutritional requirement, causing a higher uptake of ligands like lipids as seen by increased uptake of lipid binding ligands CTxB. Further, these polymeric structures gel perfectly well with scaffolds from ECM like collagen and this hybrid composition presents better ex-vivo environment to cells resulting into increased cell invasion in 3D spheroids. Taken together, this study establishes the launch pad of further exploring these DNA based hydrogels either individually or in combination with other biocomponents and together, these hydrogels are poised with attractive properties and have potential for real-world applications, including tissue engineering, drug release, cell therapy, biosensing, to name a few.

## 4. Materials and Methods

All oligonucleotides (**SI Table 1**), moviol, Triton-X, loading dye, Alexa Fluor (phalloidin) 488, moviol, poly l lysine, membrane glycoproteins (CD98 and CD147), transferrin 488, galectin, Hoechost (DAPI), Cholera toxin B-subunit (CTxB), were purchased from sigma Aldrich. Ammonium persulfate, ethidium bromide, triton X, Tetramethylethylenediamine (TEMED), Paraformaldehyde and cell culture dishes for adherent cells (treated surface) were procured from Himedia. Dulbecco’s modified Eagle’s medium (DMEM), fetal bovine serum (FBS), penicillin−streptomycin, trypsin− Ethylenediaminetetraacetic acid (EDTA) (0.25%), Collagen1 rat tail were purchased from Gibco. Tris-acetate-EDTA (TAE), Acrylamide/Bisacrylamide sol 30% (29:1 ratio for SDS PAGE) were procured from GeNei. Magnesium chloride, NaCl, KCl, Na_2_HPO_4_, and KH_2_PO_4_ were purchased from SRL, India and Santa cruez biotech respectively.

### 4.1 Cell Culture

RPE1 cells (Human retinal pigment epithelial-1 cell line) and human breast adenocarcinoma (MDA-MB-231) cells were obtained as gift from Prof. Ludger Johannes at Curie Institut, Paris, France. RPE1 and MDA-MB-231 cells were cultured in DMEM supplemented with 10% FBS and 100 IU/mL penicillin-streptomycin. For all the studies, PBS of 1× strength with pH 7.4 was used.

### 4.2 Synthesis of DNA hydrogels

Monomeric junctions of DNA ss-strands were assembled from single stranded (ssDNAs) and and each strand had a sticky end served as building unit. DNA strands of X-shaped (X1, X2, X3 and X4), Y-shaped (Y1, Y2 and Y3), T-shaped (T1, T2 and T3) at a concentration of 100 µM were individually added to 4 µM bridging agent (MgCl_2_). The resulting mixtures were annealed at a temperature range of 95-20°C for 8h. Finally, the obtained mixtures of X, Y and T-shaped DNA hydrogels were stored at 4°C for further studies.

### 4.3 Characterization

#### 4.3.1 Electrophoretic mobility shift assay

The 10% native polyacrylamide gel electrophoresis (PAGE) was prepared by using 1×Tris-acetate-EDTA (TAE) buffer. The loading sample was prepared by mixing 1-2 μL of DNA samples, 5 μL loading buffer (5×), and 1.5 μL loading dye dye, and kept for 3 min so that the dye can integrate with DNA completely. Then the loading sample was applied onto the lane and gel was run at 90 V for 90 min in 1×TBE running buffer. The bands were stained with ethidium bromide, then scanned using a Syngene Gel Documentation system.

#### 4.3.2 Dynamic light scattering

The formation for higher molecular weight DNA hydrogel by annealing was assessed by analysing the hydrodynamic diameter using Malvern Panalytical Zetasizer Nano ZS instrument. Samples containing equimolar ratio of the ssDNA oligos for respective hydrogels were analysed in low volume quartz cuvette. The hydrodynamic diameter was observed at different temperatures in decreasing order from 90°C to 5°C. The temperature difference between two steps was kept 15°C and samples were allowed to incubate at each step for 15 minutes.

#### 4.3.3 Atomic force microscopy studies

The 3D morphological characterization of synthesised DNA hydrogel was done using Bruker, multimode 8 Atomic force microscope (AFM) in tapping mode to simultaneously collect height and phase data. The hydrogel (100 μM) was drop casted on mica sheet and freeze dried before analysis. AFM scanning was done via peak force tapping mode to analyse the surface thickness, height and roughness.

#### 4.3.4 Scanning electron microscopy analysis

The porous morphology of the hydrogel was characterized using Jeol JSM-7600F Field emission scanning electron microscope (FE-SEM) at an acceleration voltage of 5 kV. To prepare samples for imaging, 10 μL of hydrogel was drop casted on carbon coated stub and freeze dried. A thin layer of platinum metal was sputter-coated on the lyophilized sample prior to imaging.

#### 4.3.5 Confocal microscopy analysis

For confocal microscopic studies, DNA hydrogel was mixed with doxorubicin (DOX) in an equimolar ratio. An appropriate amount of sample was loaded onto glass sides and freeze dried. The sample was imaged under confocal microscope in bright field as well as at 560 nm on excitation of 488 nm. The imaging was done using confocal laser scanning microscopy (Leica TCS SP8).

#### 4.3.6 Simulation studies

Conventional all-atom molecular dynamics (MD) simulation is limited to small length and time scale due to computational demand and is not suitable yet for studying gelation. So, we have used coarse-grained (CG) approach to probe the gelation of DNA nanostars. In recent years, many CG models of different resolutions have been developed to study DNA. Most of them are computationally expensive and are also too detailed, often not required to understand gelation. In this study, we have employed a minimal CG model of DNA developed by Frenkel and co-workers^32^, based on the experimental data of Xing et al.^13^ The initial all-atom models of DNA nanostars are converted to a minimal CG model comprised of beads and springs. In this model, the double-stranded arm of the DNA nanostars is represented by a large bead that interacts with other double-stranded parts with repulsive WCA potential. The single-stranded DNA ends of each nanostar’s arm were represented by small sticky beads, which attracts other single-stranded DNA ends by 12-6 Lennard Jones (LJ) potential. Each arm is represented by two large beads and a small sticky bead, and they are linked to each other by a central bead. The beads were connected by harmonic bond and angle potentials. All the force-field and simulation parameters were represented by reduced LJ units.

After building the initial CG model, we put 250 nanostars of a particular kind into a cubic box of length 30 *σ*_*LJ*_ × 30 *σ*_*LJ*_ × 30 *σ*_*LJ*_. The number density of each system is .9.25 ⨯ 10-3 The volume fractions of Y-shaped, T-shaped, and X-shaped nanostar system are 6.75,6.75,9.05, respectively. After building the initial model, each system is subjected long MD simulation using Langevin equation of motion. We cooled the system from a very high temperature (0.775 *∈*_*LJ*_*/K*_*B*_) to a very low temperature (0.075 *∈*_*LJ*_*/K*_*B*_) with an interval of 0.025 *∈*_*LJ*_*/K*_*B*_. At each temperature, the systems were subjected to long MD simulation (10^8 steps) using an integration time step of 0.005 δt in NVT ensemble. To confirm that the systems were equilibrated at each temperature, we ensured that all the thermodynamic quantities attained steady state. The damping parameter of the Langevin dynamics is chosen to be large (∼100) to mimic the overdamped condition.

### 4.4 In vitro cell studies

#### 4.4.1 Confocal imaging studies

The studies were performed on 10 mm glass cover slips coated with/without poly-l-lysine (PLL) placed in 24 well plates. Four different concentrations viz., 20, 30, 50 and 100 µM were mounted on the cover slip and incubated at 37°C for 30 min. Afterwards, trypsinized cells (approximately 40,000 cells/well) were seeded onto hydrogel treated cover slips and incubated for 24h at 37°C in 5% CO_2_. The cells were washed with PBS and fixed using 4% PFA at 37°C for 15 min. The cells were washed twice with PBS and actin filament staining was done using phalloidin 488 to check cell migration in presence of hydrogel. Cells were first treated with triton-x for 10 min at room temperature and resuspended in 500 μL phalloidin for 20 min at 37°C. After incubation, cells were washed with PBS thrice and stained with DAPI and mounted using mounting medium (moviol). All cellular fluorescent images were collected on a Leica TCS SP5 confocal microscope with a 63× oil immersion objective. A 408 nm argon laser was the excitation source for DAPI while, 488 nm argon laser was used as excitation source for phalloidin.

#### 4.4.2 Immunofluorescence assay

For this assay, coverslips placed in 24 well plates were coated with 30 and 50 µM of HGX and incubated at 37°C for 30 min. Further, trypsinized cells were seeded on to these coverslips with cell density of 40,000 cells/well and incubated at 37°C for 24 h. After 24 h, the cells were washed thrice and incubated with primary antibodies namely CD98, CD147 for 30 min at 37°C. The antibodies were prepared in serum free media in the ratio of 1:100. Fixation of cells was done using 4% PFA by incubating for 15 min at 37°C. Excess PFA was washed off with 1X PBS. The fixed cells were then permeabilized using 0.1% Triton-X-100 in PBS for 10 min at room temperature. After the removal of excess Triton-X-100, the permeabilized cells were stained with phalloidin and incubated 37°C for 20 min. After the removal of excess phalloidin was done using PBS, followed by incubation with secondary antibody prepared in 0.1% triton-x (1:250), namely, Alexa 647 in case of CD98 and CD147. Further, cells were incubated at room temperature in dark condition for 2h. Excess of secondary antibodies were removed by washing the cells with PBS thrice. Finally, the cells were mounted in a DAPI containing mounting media and stored at 4°C till imaging.

#### 4.4.3 Endocytosis pathways assessment in RPE1 cells grown over DNA HGX coated/uncoated surface

Human retinal pigment epithelial-1 cells were grown over the DNA HGX coated and uncoated glass coverslips to assess the change in cellular uptake capacity using standard endocytic probes like galectin, transferrin, and CTxB at 5 µg/mL working concentration. The cells were treated with these probes for 15 minutes at 37 ^°^C, followed by 4% PFA fixation. The fixed coverslips were then mounted on glass slides for capturing the images at 63x resolution using a confocal laser scanning microscopy (Leica TCS SP8). The raw intensity density of fluorescence images was quantified using ImageJ software and normalized with respective -HGX control. The statistic difference was performed using an unpaired t-test and indicated by an asterisk (*) in the normalized intensity plot, where **** indicates P < 0.0001; *** indicates P < 0.0003; and ** indicates P = 0.002 and ns indicates no significant difference in the data sets.

#### 4.4.4 MDA-MB 3D model preparation

MDA-MB-231 3D spheroids used for the cell spreading studies were prepared by the hanging drop culture method following a literature report with some modifications^16^. For the preparation of hanging drops, a T25 flask with 85% cellular confluency was trypsinized using 1 mL trypsin. Appropriate amount of fresh complete medium was added to the trypsinized solution and mixed well to get a homogeneous suspension. Briefly, 50 μl complete medium containing 40,000 cells were seeded in the form of drops over the inner surface of the Petri dish lid. Around 30 ml PBS was added to the Petri dish base for providing moisture environment for spheroid growth. The petri dish lid containing cell droplets was placed upside down covering the base of petriplate. The cells were allowed to grow until spheroid formation was visibly observed as the formation of cell aggregates. Spheroid formation was confirmed by visualizing cell aggregates under upright bright field optical microscope. Once the spheroids were formed, these were transferred using collagen (3mg/ml) and media in a ratio of 3:1 to 24 well plates containing 25, 50 and 100 µM HGX in required wells and incubate for ∼40 min at 37°C. Finally, the spheroids were resuspended in complete media for further incubation of 24h with and without HGX. Finally, the fixation and mounting of coverslips was done by following the same protocol as mentioned in confocal imaging studies section. The spheroids were imaged under 10x resolution.

## Acknowledgements

We sincerely thank all the members of DB group for critically reading the manuscript and their valuable feedback. SW, VM, AG and AT thank IITGN-MHRD, GoI for postdoctoral and PhD fellowship. SN acknowledges SRF fellowship from CSIR, India. DB thanks SERB, GoI for Ramanujan Fellowship, IITGN, for the startup grant, and DBT-EMR, Gujcost-DST, GSBTM and BRNS-BARC for research grants. Imaging facilities of CIF at IIT Gandhinagar are acknowledged. We sincerely thank IISc Bangalore for the computational support. The work in host labs is funded by MHRD and DST-SERB and DBT, GoI.

## Author Contributions

DB conceived the idea and planned the experiments. SW did the synthesis, characterization by confocal, and cellular area change experiments along with immunofluorescence experiments. VM did the characterization by gel electrophoresis, DLS and AFM imaging of the hydrogels, AG did the uptake experiments of Tf, Gal3 and CTxB, AT did the confocal imaging of hydrogels and 3D spheroid invasion assays, SN did the MD simulations of the hydrogels. Professors PM, CG, SD provided detailed mentorship at every step and aspect of the manuscript. SW, VM, AG, AT and SN analysed all the data and it was cross-checked by all mentors. All the authors contributed in writing and reading the manuscript draft.

## Supplementary Information

**Figure SI 1:**
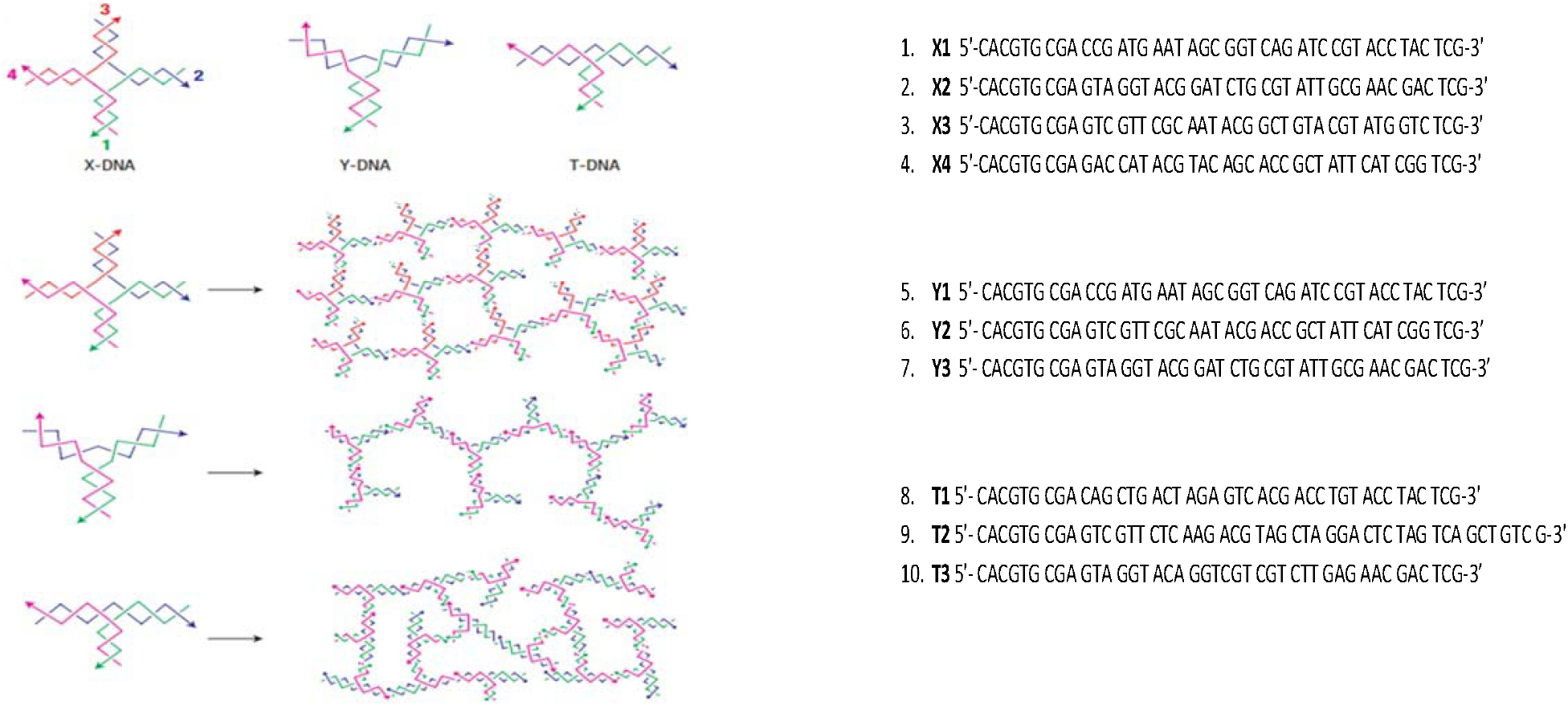
Structure and sequences of oligonucleotides used for the synthesis of DNA hydrogels. All the oligos mentioned on the left were mixed in 1:1 ratio each in PBS and Mg^2+^ and annealed in PCR machine, as described in methods section to lead to formation of respective hydrogel.

**Figure SI 2:**
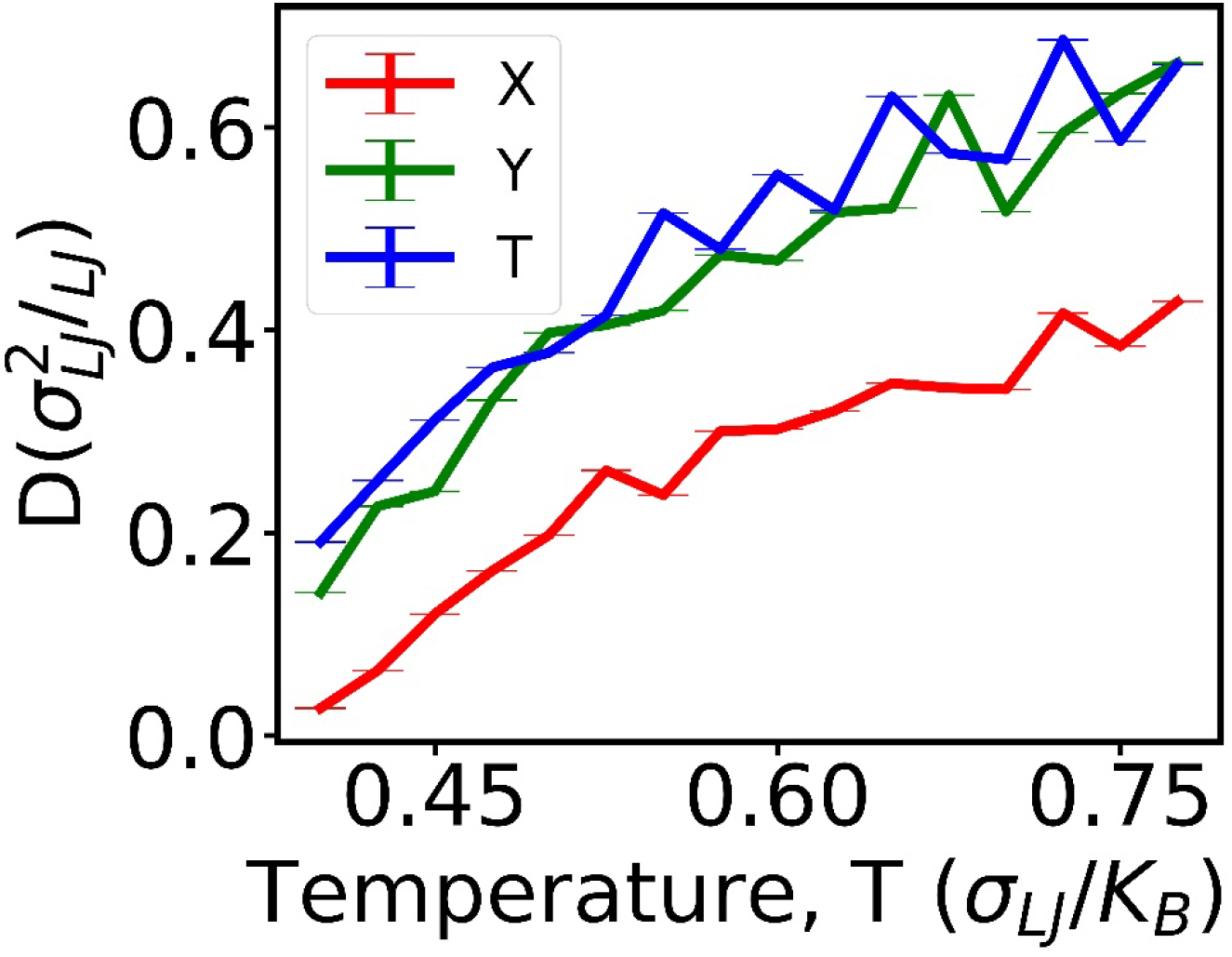
Diffusion constant as a function of temperature for different nanostar system calculated from the slope of mean-squared displacement vs time data given in the main text. The lines are guide to the eye only.

**Figure SI 3:**
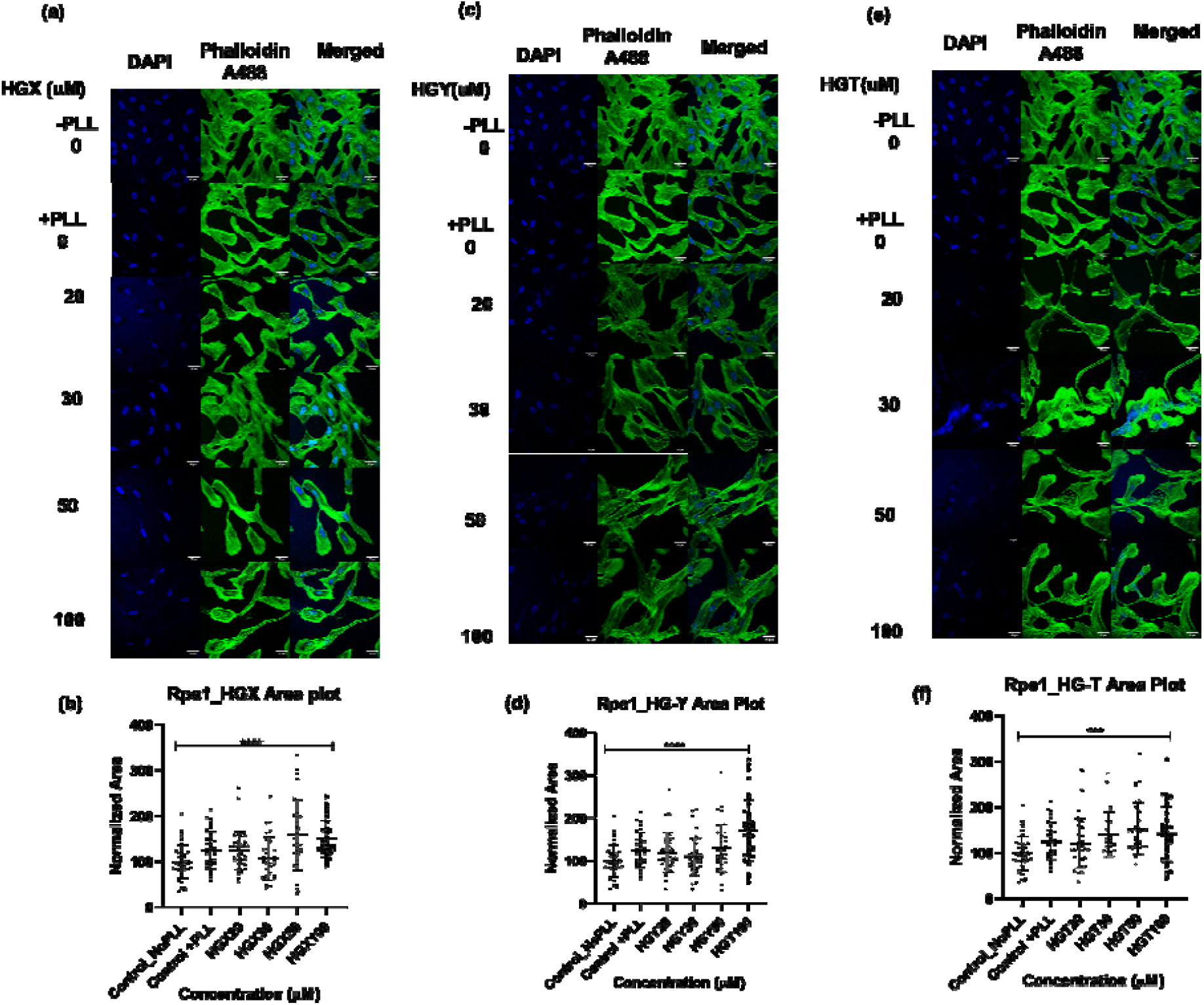
DNA hydrogels stimulate cell adhesion and enhance cell spreading. RPE cells were incubated with 20, 30 50 and 100 µM of DNA hydrogels for 24 h and the actin cytoskeleton was stained using Phalloidin. (a) HGX (c) HGY and (e) HGT. Quantification of normalized cell area w.r.t concentration. (b) HGX (d) HGY and (f) HGT. **** Indicates statistically significant value of p < 0.0001 (one-way ordinary ANOVA).

## References

(1) Yue, B. Biology of the Extracellular Matrix: An Overview. J. Glaucoma 2014, S20–S23. https://doi.org/10.1097/IJG.0000000000000108.

(2) Hinderer, S.; Layland, S. L.; Schenke-Layland, K. ECM and ECM-like Materials — Biomaterials for Applications in Regenerative Medicine and Cancer Therapy. Adv. Drug Deliv. Rev. 2016, 97, 260–269. https://doi.org/10.1016/j.addr.2015.11.019.

(3) Finke, A.; Schneider, A.; Spreng, A.; Leist, M.; Niemeyer, C. M.; Marx, A. Functionalized DNA Hydrogels Produced by Polymerase-Catalyzed Incorporation of Non-Natural Nucleotides as a Surface Coating for Cell Culture Applications. Adv. Healthc. Mater. 2019, 8 (9), 1900080. https://doi.org/10.1002/adhm.201900080.

(4) Bracaglia, L. G.; Fisher, J. P. ECM-Based Biohybrid Materials for Engineering Compliant, Matrix-Dense Tissues. Adv. Healthc. Mater. 2015, 4 (16), 2475–2487. https://doi.org/10.1002/adhm.201500236.

(5) Geckil, H.; Xu, F.; Zhang, X.; Moon, S.; Demirci, U. Engineering Hydrogels as Extracellular Matrix Mimics. Nanomed. 2010, 5 (3), 469–484. https://doi.org/10.2217/nnm.10.12.

(6) Zhong, G.; Yao, J.; Huang, X.; Luo, Y.; Wang, M.; Han, J.; Chen, F.; Yu, Y. Injectable ECM Hydrogel for Delivery of BMSCs Enabled Full-Thickness Meniscus Repair in an Orthotopic Rat Model. Bioact. Mater. 2020, 5 (4), 871–879. https://doi.org/10.1016/j.bioactmat.2020.06.008.

(7) Wang, D.; Hu, Y.; Liu, P.; Luo, D. Bioresponsive DNA Hydrogels: Beyond the Conventional Stimuli Responsiveness. Acc. Chem. Res. 2017, 50 (4), 733–739. https://doi.org/10.1021/acs.accounts.6b00581.

(8) Chen, J.; Zhu, Y.; Liu, H.; Wang, L. Tailoring DNA Self-Assembly to Build Hydrogels. Top. Curr. Chem. 2020, 378 (2), 32. https://doi.org/10.1007/s41061-020-0295-7.

(9) Sharma, A.; Sharma, P.; Roy, S. Elastin-Inspired Supramolecular Hydrogels: A Multifaceted Extracellular Matrix Protein in Biomedical Engineering. Soft Matter 2021, 17 (12), 3266–3290. https://doi.org/10.1039/D0SM02202K.

(10) Black, L. D.; Allen, P. G.; Morris, S. M.; Stone, P. J.; Suki, B. Mechanical and Failure Properties of Extracellular Matrix Sheets as a Function of Structural Protein Composition. Biophys. J. 2008, 94 (5), 1916–1929. https://doi.org/10.1529/biophysj.107.107144.

(11) Watson, J. D.; Crick, F. H. C. Molecular Structure of Nucleic Acids: A Structure for Deoxyribose Nucleic Acid. Nature 1974, 248 (5451), 765–765. https://doi.org/10.1038/248765a0.

(12) Seeman, N. C. Nucleic Acid Junctions and Lattices. J. Theor. Biol. 1982, 99 (2), 237– 247. https://doi.org/10.1016/0022-5193(82)90002-9.

(13) Xing, Z.; Caciagli, A.; Cao, T.; Stoev, I.; Zupkauskas, M.; O’Neill, T.; Wenzel, T.; Lamboll, R.; Liu, D.; Eiser, E. Microrheology of DNA Hydrogels. Proc. Natl. Acad. Sci. 2018, 115 (32), 8137–8142. https://doi.org/10.1073/pnas.1722206115.

(14) Morya, V.; Walia, S.; Mandal, B. B.; Ghoroi, C.; Bhatia, D. Functional DNA Based Hydrogels: Development, Properties and Biological Applications. ACS Biomater. Sci. Eng. 2020, 6 (11), 6021–6035. https://doi.org/10.1021/acsbiomaterials.0c01125.

(15) Xiao, M.; Lai, W.; Wang, X.; Qu, X.; Li, L.; Pei, H. DNA Mediated Self-Assembly of Multicellular Microtissues. Microphysiological Syst. 2018, 2 (1).

(16) Foty, R. A Simple Hanging Drop Cell Culture Protocol for Generation of 3D Spheroids. J. Vis. Exp. JoVE 2011, No. 51. https://doi.org/10.3791/2720.

(17) Um, S. H.; Lee, J. B.; Park, N.; Kwon, S. Y.; Umbach, C. C.; Luo, D. Enzyme-Catalysed Assembly of DNA Hydrogel. Nat. Mater. 2006, 5 (10), 797–801. https://doi.org/10.1038/nmat1741.

(18) Khalili, A. A.; Ahmad, M. R. A Review of Cell Adhesion Studies for Biomedical and Biological Applications. Int. J. Mol. Sci. 2015, 16 (8), 18149–18184. https://doi.org/10.3390/ijms160818149.

(19) Tan, J.; Gemeinhart, R. A.; Ma, M.; Mark Saltzman, W. Improved Cell Adhesion and Proliferation on Synthetic Phosphonic Acid-Containing Hydrogels. Biomaterials 2005, 26 (17), 3663–3671. https://doi.org/10.1016/j.biomaterials.2004.09.053.

(20) Chen, N.; Zhang, Z.; Soontornworajit, B.; Zhou, J.; Wang, Y. Cell Adhesion on an Artificial Extracellular Matrix Using Aptamer-Functionalized PEG Hydrogels. Biomaterials 2012, 33 (5), 1353–1362. https://doi.org/10.1016/j.biomaterials.2011.10.062.

(21) Wang, S.; Sarwat, M.; Wang, P.; Surrao, D. C.; Harkin, D. G.; John, J. A. S.; Bolle, E. C. L.; Forget, A.; Dalton, P. D.; Dargaville, T. R. Hydrogels with Cell Adhesion Peptide-Decorated Channel Walls for Cell Guidance. Macromol. Rapid Commun. 2020, 41 (15), 2000295. https://doi.org/10.1002/marc.202000295.

(22) Tang, D. D.; Gerlach, B. D. The Roles and Regulation of the Actin Cytoskeleton, Intermediate Filaments and Microtubules in Smooth Muscle Cell Migration. Respir. Res. 2017, 18 (1), 54. https://doi.org/10.1186/s12931-017-0544-7.

(23) Cantor, J. M.; Ginsberg, M. H. CD98 at the Crossroads of Adaptive Immunity and Cancer. J. Cell Sci. 2012, 125 (6), 1373–1382. https://doi.org/10.1242/jcs.096040.

(24) Iacono, K. T.; Brown, A. L.; Greene, M. I.; Saouaf, S. J. CD147 Immunoglobulin Superfamily Receptor Function and Role in Pathology. Exp. Mol. Pathol. 2007, 83 (3), 283–295. https://doi.org/10.1016/j.yexmp.2007.08.014.

(25) Wu, B.; Cui, J.; Yang, X.-M.; Liu, Z.-Y.; Song, F.; Li, L.; Jiang, J.-L.; Chen, Z.-N. Cytoplasmic Fragment of CD147 Generated by Regulated Intramembrane Proteolysis Contributes to HCC by Promoting Autophagy. Cell Death Dis. 2017, 8 (7), e2925– e2925. https://doi.org/10.1038/cddis.2017.251.

(26) Scaltriti, M.; Baselga, J. The Epidermal Growth Factor Receptor Pathway: A Model for Targeted Therapy. Clin. Cancer Res. 2006, 12 (18), 5268–5272.

(27) McMahon, H. T.; Boucrot, E. Molecular Mechanism and Physiological Functions of Clathrin-Mediated Endocytosis. Nat. Rev. Mol. Cell Biol. 2011, 12 (8), 517–533. https://doi.org/10.1038/nrm3151.

(28) Hivare, P.; Rajwar, A.; Gupta, S.; Bhatia, D. Spatiotemporal Dynamics of Endocytic Pathways Adapted by Small DNA Nanocages in Model Neuroblastoma Cell-Derived Differentiated Neurons. ACS Appl. Bio Mater. 2021, 4 (4), 3350–3359. https://doi.org/10.1021/acsabm.0c01668.

(29) Chinnapen, D. J.-F.; Chinnapen, H.; Saslowsky, D.; Lencer, W. I. Rafting with Cholera Toxin: Endocytosis and Trafficking from Plasma Membrane to ER. FEMS Microbiol. Lett. 2007, 266 (2), 129–137. https://doi.org/10.1111/j.1574-6968.2006.00545.x.

(30) Costa, E. C.; Moreira, A. F.; de Melo-Diogo, D.; Gaspar, V. M.; Carvalho, M. P.; Correia, J. 3D Tumor Spheroids: An Overview on the Tools and Techniques Used for Their Analysis. Biotechnol. Adv. 2016, 34 (8), 1427–1441. https://doi.org/10.1016/j.biotechadv.2016.11.002.

(31) 3D tumour spheroids for the prediction of the effects of radiation and hyperthermia treatments. Scientific Reports https://www.nature.com/articles/s41598-020-58569-4 (accessed 2021-06-17).

